# Impairment of proteasome-associated deubiquitinating enzyme Uchl5/UBH-4 affects autophagy

**DOI:** 10.1101/2024.04.04.588054

**Authors:** Sweta Jha, Johanna Pispa, Carina I. Holmberg

## Abstract

The autophagy-lysosomal pathway (ALP) and the ubiquitin-proteasome system (UPS) are the two major intracellular proteolytic systems that mediate protein turnover in eukaryotes. Although a crosstalk exists between these two systems, it is still unclear how UPS and ALP interact *in vivo*. Here, we have investigated how impaired function of the proteasome-associated deubiquitinating enzyme (DUB) Uchl5/UBH-4, a regulator of proteasome activity, affects autophagy in human cells and in various tissues in a multicellular organism. We have used the established GFP-LC3-RFP-LC3ΔG autophagy reporter HeLa cell line and show that downregulation of *Uchl5* by siRNA reduces autophagy, which is similar to a previously reported role of the proteasome-associated DUB Usp14 on autophagy. Exposing *Caenorhabditis elegans* carrying the autophagy reporter mCherry::GFP::LGG-1 to *ubh-4* or *usp-14* RNAi, or to their pharmacological inhibitors, results in diverse effects regarding the numbers of autophagosomes and autolysosomes in the intestine, hypodermal seam cells and the pharynx. Our results reveal that the proteasome-associated DUBs Uchl5/UBH-4 and Usp14 affect autophagy in a differential tissue manner. A deeper insight into the interplay between UPS and ALP in various tissues *in vivo* has the potential to promote development of therapeutic approaches for disorders associated with proteostasis dysfunction.

**Summary:** Modulation of UPS via pharmacological or genetic impairment of the proteasome-associated DUB Uchl5/UBH-4 affects autophagy in human cells and in a tissue-differential manner in *C. elegans*.

## Introduction

Protein homeostasis is a dynamic balance between the production and degradation of proteins and is essential for cell survival and growth. In eukaryotes, the turnover of proteins is facilitated by two main intracellular proteolytic systems: the autophagy-lysosomal pathway (ALP) (hereafter referred to as autophagy), and the ubiquitin-proteasome system (UPS). These two proteolytic systems are essential components of the cellular protein quality control system, but also crucial for the maintenance of the amino acid pools and energy balance [reviewed in (Schreiber & Peter, 2014) (Pohl & Dikic, 2019)].

Autophagy is a highly regulated proteolytic pathway that is well conserved from yeast to humans. It is responsible for degradation of mainly long-lived proteins, cytoplasmic organelles, and other cellular components by delivering them to the lysosomes, and the breakdown products are reused for cellular processes [reviewed in (Wen & Klionsky, 2020)]. The autophagy process starts with the formation of an isolation membrane, the phagophore, which elongates to engulf the substrate(s) and forms the double-layered autophagosome. Autophagosomes then fuse with late endosomes and lysosomes, which leads to the formation of autolysosomes, where the substrate(s) is degraded by lysosomal hydrolases (Kametaka et al., 1998) (Mizushima, 2007) (Mizushima, 2018) (Sun-Wang et al., 2020). Autophagy is mediated through the conserved action of the Atg protein family, of which Atg8, a ubiquitin-like protein, is a key participant in the autophagic process in yeast. Atg8 and its mammalian homologs GABARAP and LC3 play crucial role in various stages of autophagy, encompassing initiation, cargo recognition and engulfment, as well as the closure of autophagosome (Mizushima et al., 1998) (Nakatogawa et al., 2007) (Knorr et al., 2014) (Klionsky et al., 2021). At the autophagosome membrane, Atg8/GABARAP/LC3 is conjugated with phosphatidylethanolamine (PE), and the Atg8-PE/LC3-II serves as a well-established marker for assessment of autophagy (Nakatogawa et al., 2007) [reviewed in (Klionsky et al., 2021) (Yamamoto et al., 2023)]. In a multicellular organism, a role of autophagy was firstly reported in *Caenorhabditis elegans* dauer development (Meléndez et al., 2003) and autophagy has since been shown to be essential also for embryogenesis, longevity, and stress responses (Zhao et al., 2009) (Tian et al., 2010) (Palmisano & Meléndez, 2019) (Meléndez et al., 2003) (Alberti et al., 2010) (Wu et al., 2015) (Chang et al., 2017). *C. elegans* has two Atg8 homologs, i.e., LGG-1 and LGG-2 corresponding to GABARAP and LC3, respectively. Both LGG-1 and LGG-2 localize to autophagosomes and LGG-1 is required for the recruitment of LGG-2 (Alberti et al., 2010) (Manil-Ségalen et al., 2014). The structural conservation of LGG-1/GABARAP and LGG-2/LC3 emphasizes their pivotal role in autophagy during developmental stages, longevity, and responses to stress (Meléndez et al., 2003) (Alberti et al., 2010) (Wu et al., 2015) (Chang et al., 2017). Recently, a lipidation-independent role of LGG-1 in autophagy was reported (Leboutet et al., 2023).

UPS is the primary proteolytic pathway responsible for degradation of soluble, short-lived, and misfolded proteins in the cytosol and nucleus. UPS-mediated proteolytic degradation is a multistep process, where polyubiquitination of proteasomal substrates occurs through the action of a cascade of three different classes of enzymes: ubiquitin-activating enzymes (E1), ubiquitin-conjugating enzymes (E2) and ubiquitin ligases (E3). The polyubiquitinated substrates are then degraded by the evolutionarily conserved 26S proteasome, a large ATP-dependent multicatalytic protease complex (J. Chen et al., 2020) [reviewed in (Finley, 2009) (Dikic, 2017) (Raffeiner et al., 2023)]. Prior to degradation ubiquitin chains are removed from the substrates by three proteasome-associated deubiquitinating enzymes (DUBs); the cysteine proteases UchL5/Uch37 (Stone et al., 2004) and Usp14 (Borodovsky et al., 2001), and the metalloprotease Rpn11 (Verma et al., 2002). Of these three proteasome-associated DUBs, Rpn-11 is a subunit of the proteasome, whereas Uchl5 and Usp14 bind to the proteasome [reviewed in (Collins & Goldberg, 2017; Finley, 2009) (Kocaturk & Gozuacik, 2018)]. Rpn11 is responsible for en bloc removal of ubiquitin chains. Usp14 functions either by trimming or by en bloc removal of ubiquitin chains, and Uchl5 functions by trimming or by debranching polyubiquitin chains (Hanna et al., 2006) (Finley, 2009) (M. J. Lee et al., 2011), (B.-H. Lee et al., 2016) (Deol et al., 2020). Due to their modes of action Usp14 and Uchl5 affect the kinetics and affinity of the substrate-proteasome interaction, thereby influencing the proteolytic capacity of the proteasome. Impaired Usp14 or Uchl5 function have been reported to enhance proteasomal degradation of substrates (Lam et al., 1997) (Hanna et al., 2006) (Koulich et al., 2008) (Matilainen et al., 2013) (H. T. Kim & Goldberg, 2017) (H. T. Kim & Goldberg, 2018) (J. H. Lee et al., 2018) (Deol et al., 2020), but also to cause accumulation of proteasomal substrates (J. H. Lee et al., 2018) (Chadchankar et al., 2019)

In the past, autophagy and UPS were believed to function as independent systems targeting distinct substrates, however, accumulating evidence describes interactions between these two systems including some shared substrates [reviewed in (Pohl & Dikic, 2019) (Raffeiner et al., 2023)]. The interplay is not always a direct compensatory mechanism but exhibits complexity and diversity. Studies on pharmacological and genetic impairment of autophagy have reported contrasting outcomes, as both inhibition of proteasomal degradation as well as upregulation of proteasome activity have been reported. Ding et al. have shown that siRNA knockdown of Atg6/Beclin1 or Atg8/LC3 results in accumulation of polyubiquitinated proteins in HCT116 human colorectal carcinoma cells (Ding et al., 2007). Additionally, impairment of autophagy by siRNA or chemical treatments in HeLa cervical cancer cells or by Atg5 knockout in MEF (Mouse Embryonic Fibroblasts) cells induces accumulation of a UPS reporter due to decreased proteasomal degradation (Korolchuk et al., 2009). In neuroblastoma cells, inhibition of lysosomal function has been reported to decreased proteasome activity (Qiao & Zhang, 2009). However, there are also studies showing that autophagy modulation has an opposite effect on UPS. Wang et al. have demonstrated that inhibition of autophagy via chemical treatments or downregulation of Atg5 or Atg7 leads to upregulation of proteasome activity and increased expression of proteasomal subunits in colon cancer cell lines SW1116 and HCT116 (Wang et al., 2013). Kim et al. have shown that autophagy-defective Atg5 knockout MEF cells increased proteasome activity and starvation-induced autophagy resulted in decreased proteasome activity without affecting the stability of proteasome subunits in HEK293 human embryonic kidney cells (E. Kim et al., 2018). We have previously shown that RNAi downregulation of various autophagy genes do not result in systemic upregulation of UPS function in *C. elegans,* but depending on the target gene the outcome on UPS function or proteasome expression varies in a tissue-specific manner (Jha & Holmberg, 2020). Several studies on impaired proteasomes on the other hand, have reported an induction of autophagy in human cell lines, mice cells and in drosophila (Q. Zheng et al., 2011) (Shen et al., 2013) (Lőw et al., 2013) (Fan et al., 2016) (Li et al., 2019).

We have previously shown that Uchl5 depletion enhances proteasomal substrate degradation in human U-2 OS (osteosarcoma) cells and that the *C. elegans* homolog UBH-4 regulates proteasome activity in *C. elegans* intestine, and affects the lifespan and health span of the animal (Matilainen et al., 2013). Here, we investigated the effect of Uchl5/ UBH-4 on autophagy in human cells and *C. elegans*. We also performed comparison studies between Uchl5 and Usp14, as Usp14 downregulation has been shown to decrease autophagic flux in HEK293 and MEF cells (E. Kim et al., 2018). Our data reveal that *Uchl5 and Usp14* siRNA treatments or pharmacological inhibition reduce autophagy in GFP-LC3-RFP-LC3ΔG HeLa cells. When investigating the impact of these DUBs on autophagy at the tissue level, our results show that downregulation of *ubh-4* and *usp-14* by RNAi or by using pharmacological inhibitors causes differential autophagic responses in the intestine, hypodermal seam cells and the pharynx in *C. elegans*. Our studies highlight the complexity in the interaction between the ubiquitin-proteasome system and autophagy in a multicellular organism.

## Results

### Knockdown of deubiquitinating enzyme (DUB) *Uchl5* reduces autophagy

To investigate how the proteasome-associated deubiquitinating enzyme (DUB) Uchl5 affects autophagy, we downregulated *Uchl5* by siRNA and analyzed the effect on autophagic flux. We used the previously developed HeLa cell line expressing the autophagy reporter GFP-LC3-RFP-LC3ΔG, which is cleaved into equimolar amounts of GFP-LC3 and RFP-LC3ΔG by the endogenous protease ATG4 in the cell (Kaizuka et al., 2016). While phosphatidylethanolamine-conjugated GFP-LC3 (LC3-II) in the autophagosomes is degraded upon fusion with lysosomes, the stable RFP-LC3ΔG present in the cytosol serves as an internal control, and thus, the fluorescence ratio of the GFP/RFP signal reversely correspond to autophagic activity (Kaizuka et al., 2016) (Fig S1A).

Treatment of GFP-LC3-RFP-LC3ΔG HeLa cells with *Uchl5* siRNA resulted in reduced *Uchl5* mRNA and Uchl5 protein levels, respectively (Fig S1B and S1C). We also downregulated the proteasome-associated DUB *Usp14* (Fig S1B and S1C), the inhibition of which has previously been shown to impair autophagy at the autophagosome-lysosome fusion step in HEK293 and MEF cells (E. Kim et al., 2018), to enable comparison of autophagy studies with Uchl5. Both *Uchl5* and *Usp14* siRNA resulted in an increased number of GFP-LC3 puncta as measured by live imaging, and a significant increase in the GFP/RFP ratio was detected, indicating inhibition of autophagic activity (Fig 1A). Next, we checked the levels of LC3 in cell lysates upon *Uchl5* or *Usp14* knockdown and observed a significant accumulation of LC3-II in these cells (Fig 1B). Similarly, we detected an accumulation of the autophagy marker/substrate p62 upon *Uchl5* or *Usp14* siRNA treatment (Fig 1C). For validation, we also treated cells with Bafilomycin A (BAFA) and observed an expected increase in the GFP/RFP ratio (Fig S2A) as well as accumulation of LC3 and p62 (Fig S2B), in agreement with previous reports on inhibition of autophagy upon BAFA treatment (Redmann et al., 2017) (Kaizuka et al., 2016). Taken together, our *in vivo* and *in vitro* data show that similar to Usp14, Uchl5 reduction decreases autophagy in HeLa cells.

**Figure 1.**
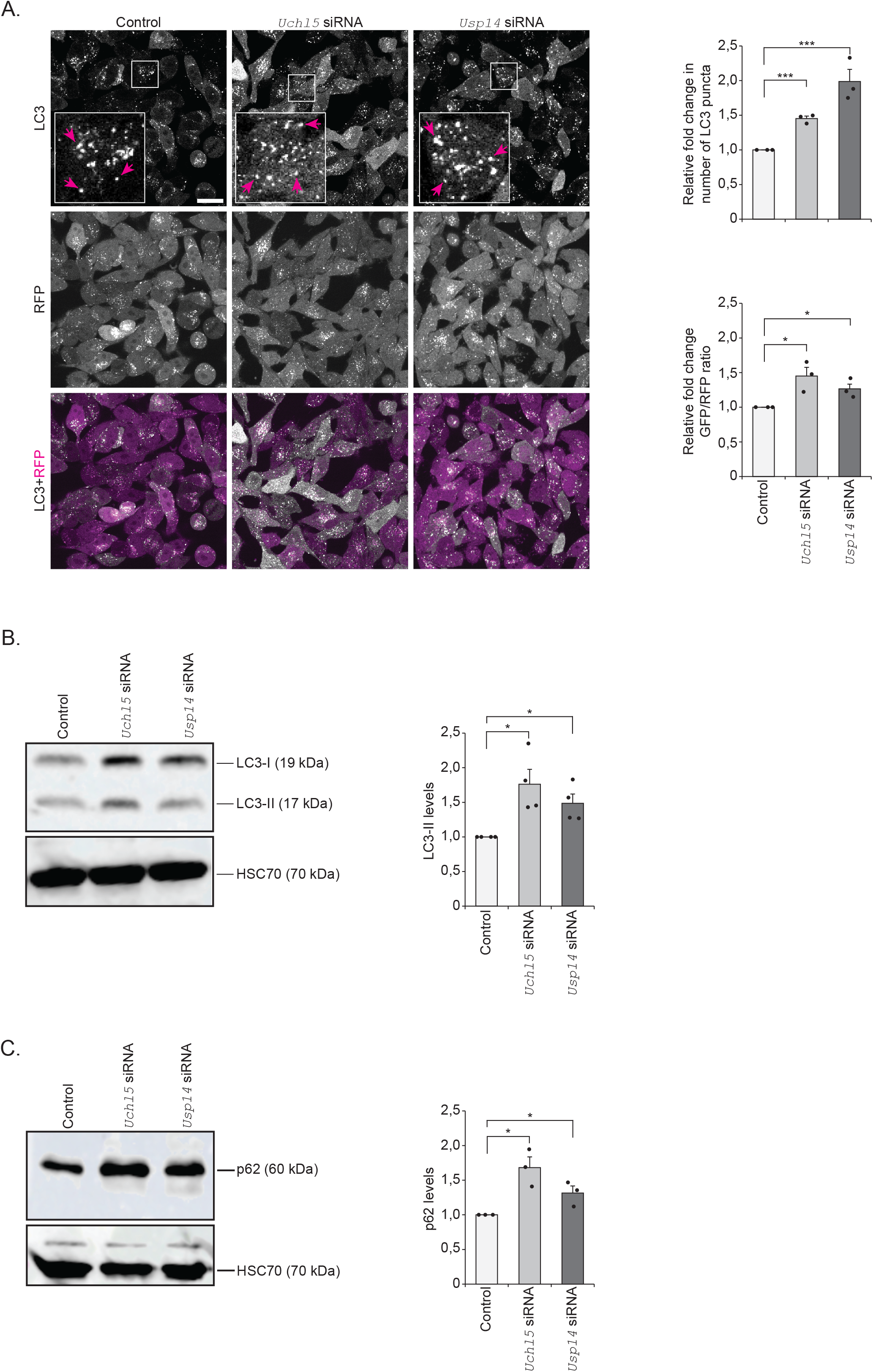
Downregulation of proteasome-associated DUBs *Uchl5* and *Usp14* reduces autophagy. **A.** Fluorescence confocal images of control, *Uchl5* or *Usp14* siRNA treated GFP-LC3-RFP-LC3ΔG HeLa cells after 48h post-transfection. Insets show enlarged view of the indicated areas. Magenta arrows point to some of the GFP-LC3 puncta. Scale bar, 20 µm. The right upper graph shows quantification of relative fold change in the number of GFP-LC3 puncta per image (Control set at 1). The right lower graph shows quantification of the relative fold change in the ratio of GFP to RFP per image (Control set at 1). Results are from three independent experiments (total 15-17 images were analyzed). Error bars, SEM, *p<0,05, ***p<0,001 compared to control. **B and C**. GFP-LC3-RFP-LC3ΔG HeLa cells treated with control, *Uchl5* or *Usp14* siRNA for 48h. Whole cell extracts were analyzed by SDS-PAGE and immunoblotted against LC3, p62 and HSC70. The graphs (on right panels) show average fold change in levels of LC3 (B) and p62 (C) normalized against HSC70. Results are the mean of quantifications from 3-4 independent experiments. Error bars, SEM, *p<0,05 compared to the control (set as 1).

### Pharmacological inhibition of proteasome-associated DUBs decreases autophagy

To complement the studies on genetic downregulation of *Uchl5* and *Usp14*, we also investigated the effect of pharmacological inhibition of these two proteasome-associated DUBs in the GFP-LC3-RFP-LC3ΔG HeLa cells. Of the commonly used DUB inhibitors, b-AP15 blocks the DUB activity of both Uchl5 and Usp14 (D’Arcy et al., 2011) and IU1 inhibits specifically Usp14 activity (B.-H. Lee et al., 2010). To our knowledge there is so far no specific inhibitor for Uchl5, and therefore, we performed comparison analysis between dual inhibition of Uchl5 and Usp14 and single inhibition of Usp14 to reveal insight on the action of Uchl5 on autophagy. We investigated the autophagy in live GFP-LC3-RFP-LC3ΔG HeLa cells treated with b-AP15 or IU1, and the number of GFP-LC3 puncta as well as the GFP/RFP ratio were analyzed. We detected that b-AP15 treatment increased the number of GFP-LC3 puncta as well as the GFP/RFP ratio (Fig 2A). The b-AP15 treatment also resulted in increased amount of LC3-II (Fig 2B) and p62 (an autophagosome substrate) (Fig 2C), as analyzed by Western blotting. These results are indicative of reduced autophagy and correlate with the reduced autophagy we detected upon *Uchl5* or *Usp14* downregulation by siRNA (Fig 1). Similarly, treatment with IU1 resulted in increased number of GFP-LC3 puncta and the GFP/RFP ratio, indicating a reduction of autophagy in GFP-LC3-RFP-LC3ΔG HeLa cells (Fig S3A). Consistent with a previous report by Kim et al 2018, we also observed a significant accumulation of both LC3-II and p62 upon IU1 treatment (E. Kim et al., 2018) (Fig S3B and S3C). Taken together, our results reveal that both downregulation and pharmacological inhibition of Uchl5, as well as Usp14, causes reduction of autophagy.

**Figure 2.**
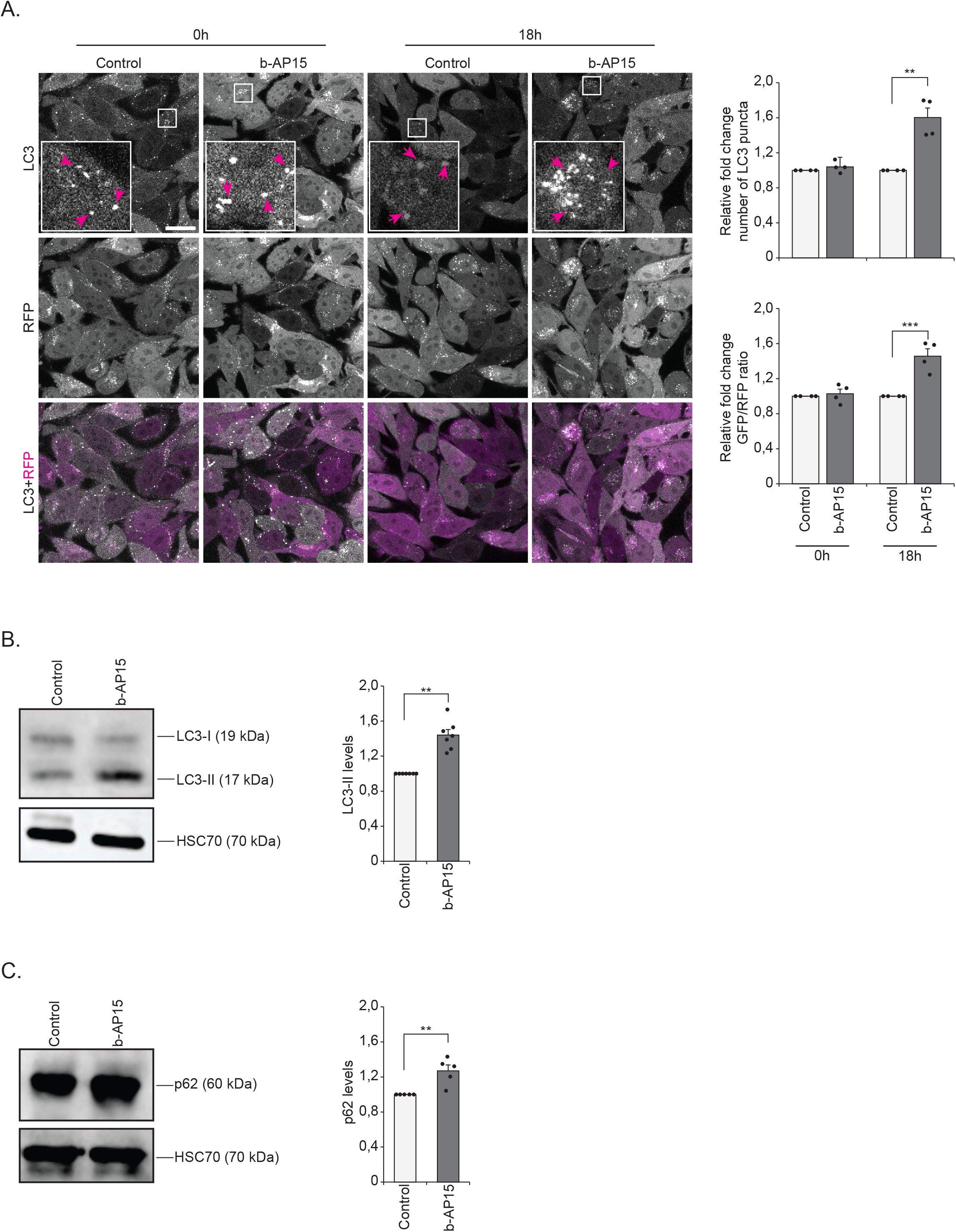
Pharmacological inhibition of proteasome-associated DUBs Uchl5 and Usp14 reduces autophagy. **A.** Fluorescence confocal images of control (DMSO) or b-AP15 (1µM) treated GFP-LC3-RFP-LC3ΔG HeLa cells after 18h of treatment. Insets show enlarged view of the indicated areas. Magenta arrows point to some of the puncta. Scale bar, 20 µm. The right upper graph shows the quantification of relative fold change in the number of GFP-LC3 puncta per image (Control set at 1). The right lower graph shows the quantification of the relative fold change in the ratio of GFP to RFP per image (Control set at 1). Results are from four independent experiments (total 15-20 images were analyzed). Error bars, SEM, **p<0,01, ***p<0,001 compared to control. **B and C**. GFP-LC3-RFP-LC3ΔG HeLa cells treated with control (DMSO) or b-AP15 (1µM) for 18h. whole cells extract were analyzed by SDS-PAGE and immunoblotted against LC3, p62 and HSC70. The graphs (on right panel) show average fold change in levels of LC3 (B) and p62 (C) normalized against HSC70. Results are the mean of quantifications from 5-7 independent experiments. Error bars, SEM, **p<0,01 compared to the control (set as 1).

### Differential autophagic tissue responses to downregulation of the proteasome-associated DUBs *ubh-4* and *usp-14* in *C. elegans*

Previously, we have reported that impairment of autophagy affects proteasome function differently in the intestine and body-wall muscle in *C. elegans* (Jha & Holmberg, 2020). To address whether downregulation of proteasome-associated DUBs *ubh-4*, the *uchl5* homolog, and *usp-14* induces a tissue-specific or systemic effect on autophagy in *C. elegans*, we downregulated *ubh-4* and *usp-14* by RNAi-feeding and analyzed autophagy in intestinal cells, hypodermal seam cells and pharynx (Fig 3A). These cell types were chosen as they have previously been used for assessing autophagy in *C. elegans* (Zhang et al., 2015) (Chang et al., 2017). The RNAi treatment resulted in efficient reduction of *ubh-4* and *usp-14* mRNA levels, as measured in whole animal lysates (Fig S4). For monitoring autophagy, we used the previously published dual-fluorescent marker strain expressing mCherry::GFP::LGG-1 driven by the endogenous *lgg-1* promoter (Chang et al., 2017) (Fig 3B). In these animals, the autophagosomes (APs) are indicated by puncta positive for both GFP and mCherry, and the autolysosomes (ALs) are visualized as red puncta (mCherry) due to lysosomal quenching of GFP (Chang et al., 2017) (Fig 3C).

**Figure 3.**
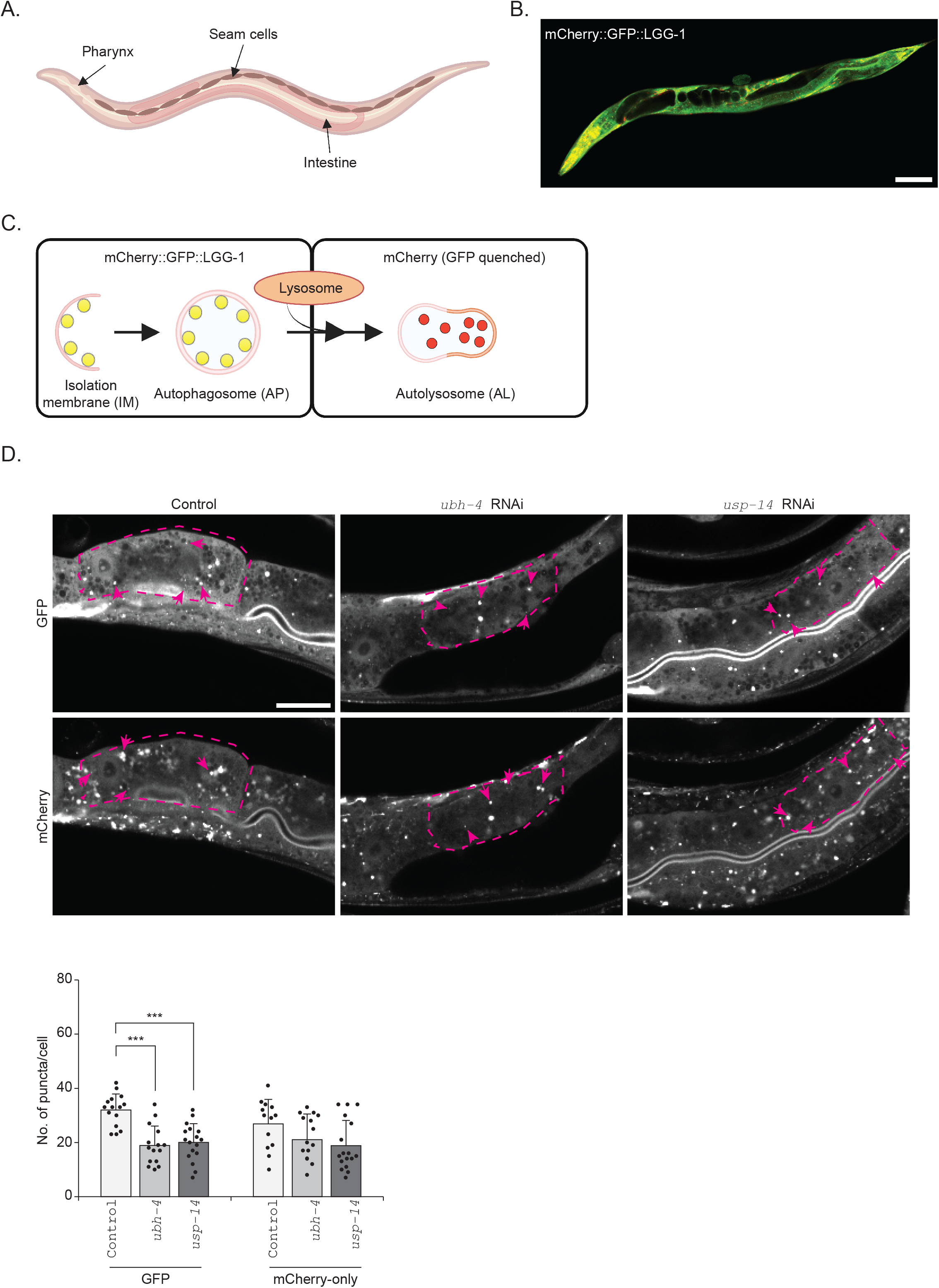
The autophagosome pool size decreases upon downregulation of the proteasome-associated DUBs *ubh-4* and *usp-14* in intestinal cells of *C. elegans*. **A.** Schematic representation of the tissues (studied in this article) of an adult *C. elegans.* B. Fluorescence image of a day 1 old animal expressing mCherry::GFP::LGG-1. Scale bar: 100 µm. Note that the image was taken with different setting for GFP and mCherry. **C.** Schematic representation of the fluorescence states of mCherry::GFP::LGG-1 at the different stages of autophagy (Isolation membrane, IM; Autophagosome, AP; Autolysosome, AL). **D.** Representative confocal micrographs of control, *ubh-4* or *usp-14* RNAi-treated mCherry::GFP::LGG-1 animals at day 1 of adulthood. Individual intestinal cells are outlined with magenta dashed lines and the magenta arrows point to some of the puncta. Scale bar: 50 µm. Note that the mCherry was imaged with lower gain setting. The graph (below) shows the quantification of the number of puncta positive for GFP and mCherry-only in individual intestinal cells. Results are from three independent experiments. Puncta were counted from a total of 15-18 individual intestinal cells from 12-15 animals (all distinct puncta of variable sizes were counted). Error bars, SEM, ***p<0,001 compared to control.

We started the RNAi feeding at L1 larval stage, imaged live animals at day 1 of adulthood, and subsequently manually counted fluorescent puncta in individual intestinal cells, hypodermal seam cells and pharynx of the animals. The AP puncta positive for both GFP and mCherry (yellow puncta) were not easily distinguishable from the diffused fluorescence, as the mCherry fluorescence signal was much brighter than the GFP even with optimized imaging settings for both channels. As the study by Chang *et al*. reported that GFP positive puncta are equivalent to puncta positive for both GFP and mCherry, we similarly counted APs as GFP puncta and ALs as mCherry-only puncta (the total number of mCherry positive puncta with subtraction of the GFP positive puncta). Our control experiments by RNAi of the well-established autophagy genes *lgg-1* (homolog of atg-8) and *rab-7* (homolog of mammalian RAB7) confirmed expected results, as knockdown of *lgg-1* profoundly decreased the appearance of both APs and ALs and *rab-7* RNAi increased APs and decreased ALs in intestinal cells and hypodermal seam cells, which is consistent with its role in the fusion of autophagosomes to lysosomes (Fig S5A). Upon *ubh-4* or *usp-14* RNAi, we observed a significant decrease in the number of APs without an effect on the number of ALs in intestinal cells (Fig 3D). Our result suggests that in intestinal cells, the early stage of autophagy, *i.e.,* AP formation, is reduced, but not the AP fusion with the lysosome, upon downregulation of *ubh-4* or *usp-14.* Similarly, the hypodermal seam cells displayed decreased number of APs, but not ALs, when the animals were exposed to *usp-*14 RNAi, (Fig 4A). Downregulation of *ubh-4* on the other hand did not affect the number of formed APs, but resulted in an increased number of ALs, suggesting slow lysosomal degradation (Fig 4A).

**Figure 4.**
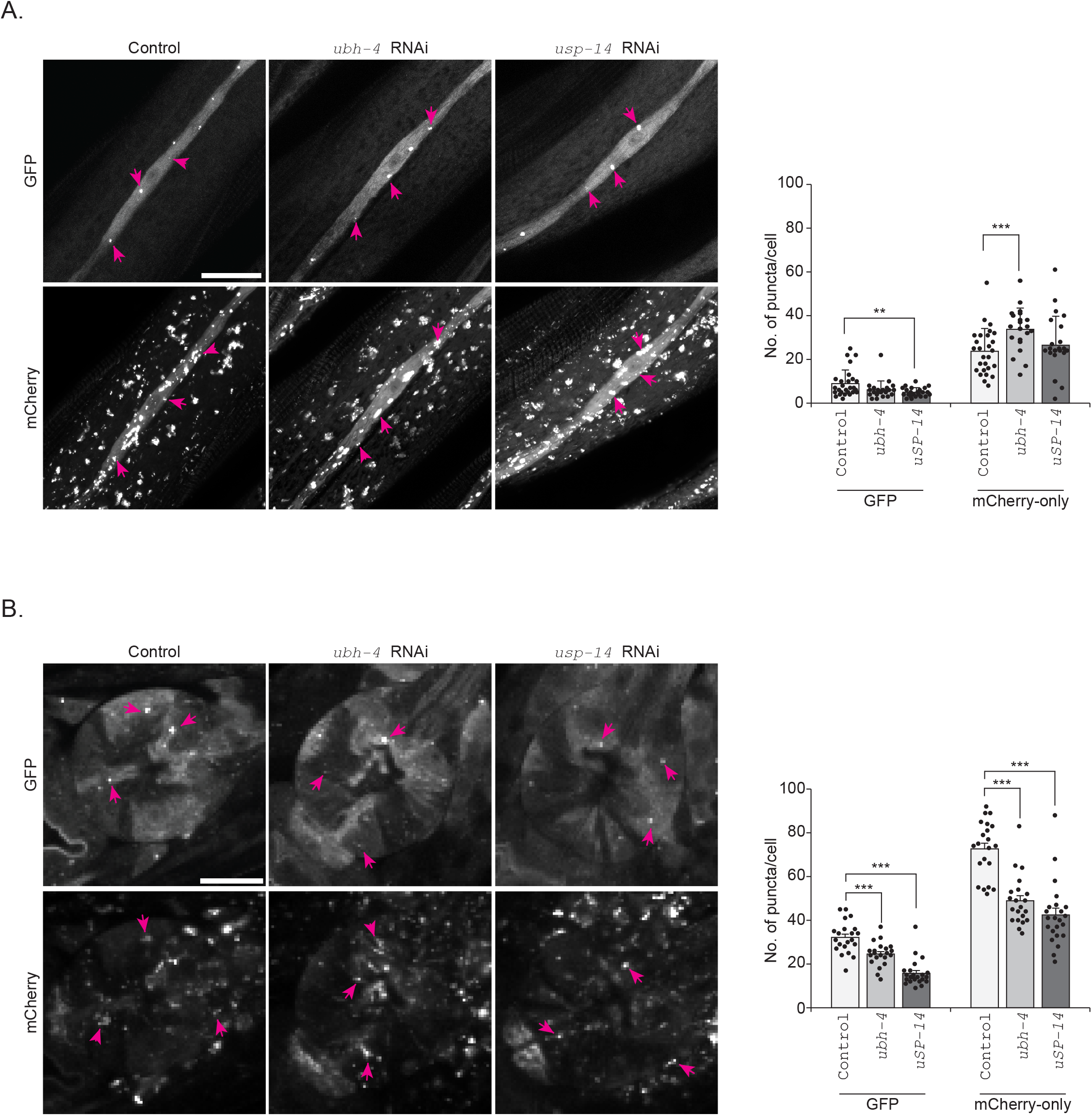
Downregulation of the proteasome-associated DUBs *ubh-4* and *usp-14* affects autophagy differently in the pharynx and in hypodermal seam cells. Fluorescence confocal micrographs of control, *ubh-4* or *usp14* RNAi-treated mCherry::GFP::LGG-1 animals showing hypodermal seam cells **(A)** and, the pharynx **(B).** Graphs show the quantification of the number of puncta positive for GFP and mCherry-only. Results are from five independent (for hypodermal seam cells) or three independent (for pharynx (second generation)) experiments. Puncta were counted from a total of 20-25 pharynges from 20-25 animals and 25-30 hypodermal seam cells from 20-25 animals. Error bars, SEM, **p<0,01 and ***p<0,001 compared to control.

We next analyzed the effect of downregulation of *ubh-4* and *usp-14* on autophagy in the pharynx. Previous studies have reported that pharynx is resistant to first generation RNAi, but sensitive in animals exposed to RNAi for the second generation (Kumsta & Hansen, 2012) (Shiu & Hunter, 2017). Accordingly, we observed no effect on the number of APs and ALs upon first generation RNAi of *lgg-1, rab-7, ubh-4* or *usp-14* (Figs S5A and S5B), whereas the effect of autophagy was clearly detected in the pharynx of animal continuously exposed to *lgg-1* or *rab-7* RNAi for two generations (Fig S5A). Animals exposed to continuous *ubh-4* or *usp-14* RNAi for two generations displayed a significant decrease in the number of APs and ALs in the pharynx (Fig 4B).

Altogether, our results reveal that *usp-14* and *ubh-4* can have similar or distinct effects on the AP and AL stages of autophagy in the intestine, hypodermal seam cells and pharynx in *C. elegans* (summarized in Fig 6).

**Figure 5.**
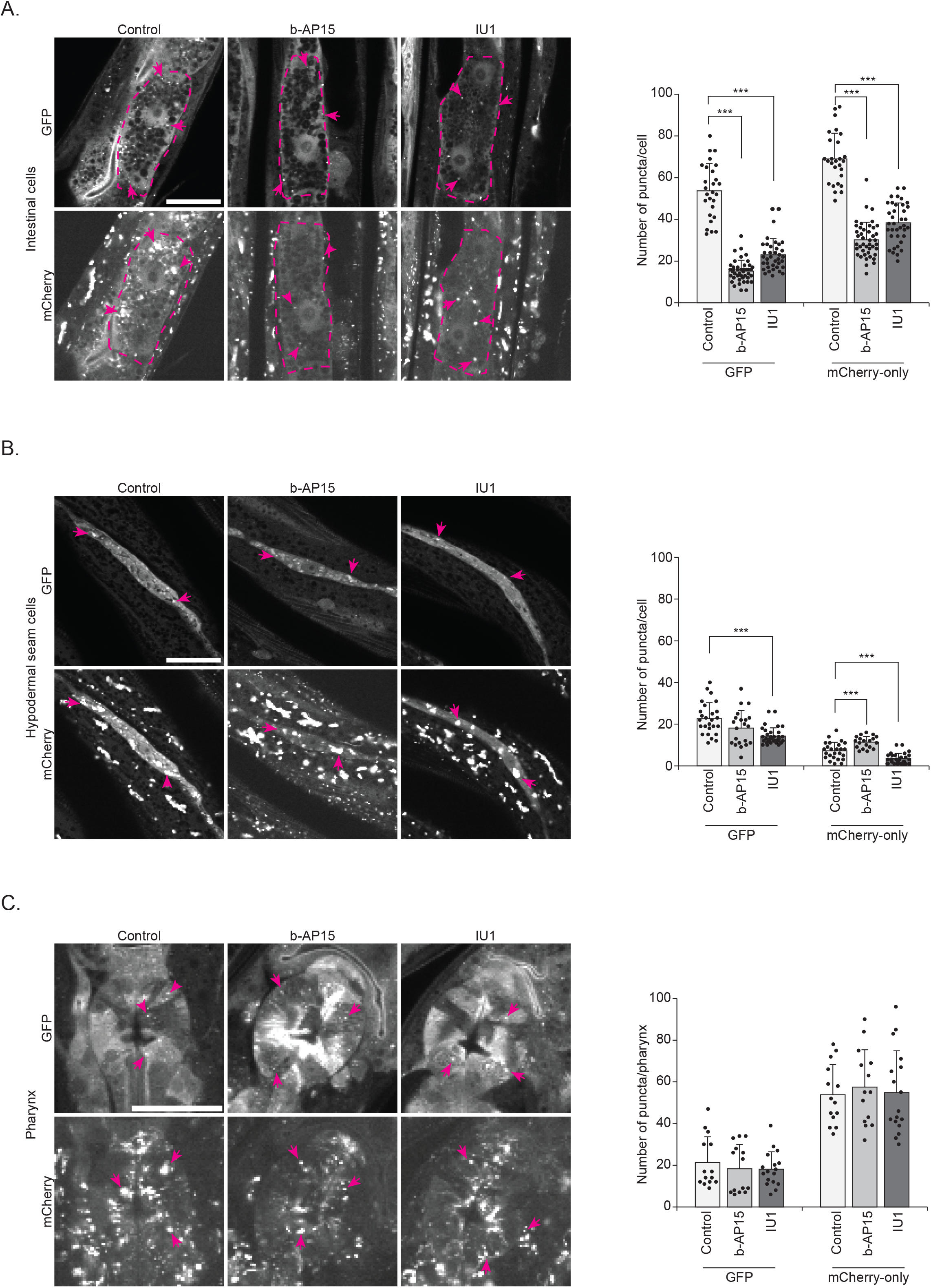
Pharmacological inhibition of the proteasome-associated DUBs *ubh-4* and *usp-14* affects autophagy differently in different tissues. Representative fluorescence confocal micrographs of control (DMSO), b-AP15 (10 µM) or IU1 (100 µM) treated mCherry::GFP::LGG-1 animals showing the intestinal cells **(A)**, hypodermal seam cells **(B),** and the pharynx **(C).** Graphs show the quantification of the number of puncta positive for GFP and mCherry-only in the corresponding cells. Results are from three independent experiments. Puncta were counted from a total of 30-35 individual intestinal cells from 20-25 animals, 30-35 hypodermal seam cells from 20-25 animals and 15-17 pharynges from 15-17 animals. Error bars, SEM, ***p<0,001 compared to control.

**Figure 6.**
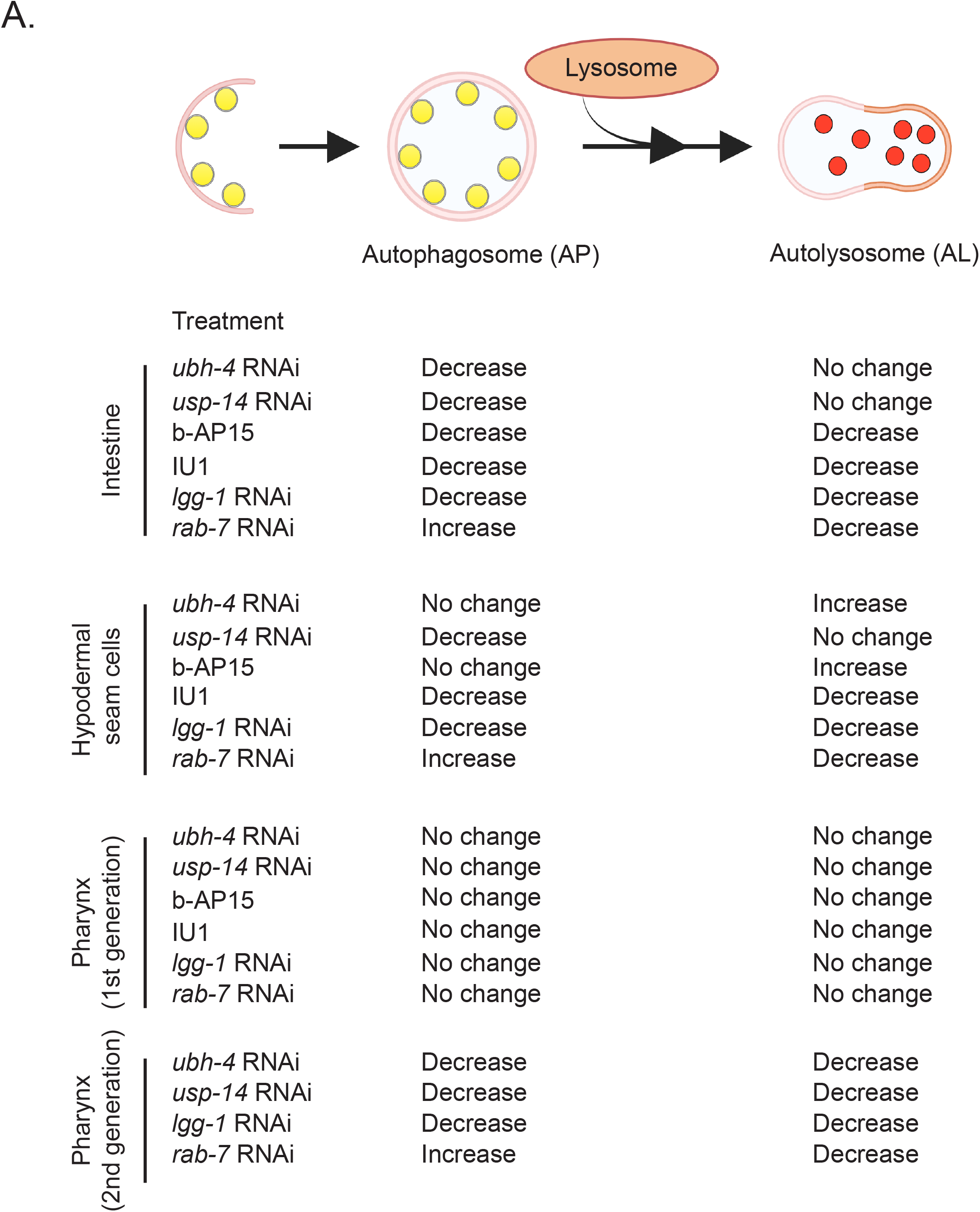
Summary of the effects of *ubh-4* and *usp-14* impairments on autophagy in the intestine, hypodermal seam cells and the pharynx.

### Pharmacological inhibitors of proteasome-associated DUBs affect autophagy in a tissue-specific manner in *C. elegans*

To complement the studies on genetic downregulation of *ubh-4* and *usp-14*, we also investigated the effect of the deubiquitinase inhibitors, b-AP15 and IU1, which are commonly used for mammalian cell culture studies, on autophagy in *C. elegans.* When animals expressing the mCherry::GFP::LGG-1 reporter were exposed to b-AP15 or IU1 from the L1 stage to analysis at day 1 of adulthood, we detected that both inhibitor treatments decreased the number of APs as well as ALs in the intestine (Fig 5A). Our data reveals that pharmacological inhibitor treatment of both *ubh-4* and *usp-14* suppresses autophagy at early stage.

In hypodermal seam cells, b-AP15 treatment did not affect the number of APs but resulted in more ALs (Fig 5B) suggesting a faster fusion of APs with lysosomes. The suppression of *usp-14* using IU1 decreased both APs and ALs (Fig 5B) in hypodermal seam cells, indicating that a reduction in *usp-14* affects the early stage of autophagy. Notable, treatment with the dual inhibitor b-AP15 elicited an opposite response in terms of the effect on number of ALs. Treatments with b-AP15 and IU1 did not affect the number of APs or ALs in the pharynx (Fig 5C).

As the inhibitors b-AP15 and IU1 affect the proteasome-associated DUBs (B.-H. Lee et al., 2010), we would expect to detect a change in the accumulation of proteasomal substrates in *C. elegans*. We, therefore, used these inhibitors on our previously established fluorescent polyubiquitin reporter strain, which measures endogenous Lys-48-linked polyubiquitinated proteasomal substrates in the intestine (Matilainen et al., 2013) (Matilainen et al., 2016). The polyubiquitin reporter animals were treated with b-AP15 or IU1 from the L1 larval stage until day 1 of adulthood. We detected an increase in fluorescence upon b-AP15 treatment reflecting increased accumulation of polyubiquitinated proteins in the intestine (Figs S6A and S6B). IU1 treatment on the other hand resulted in decreased amount of polyubiquitinated proteins in the intestine (Fig S6A and 6B). Our *C. elegans* results support previous reports on the effect of b-AP15 and IU1 on proteasomal substrates in mammalian cells (D’Arcy et al., 2011) (B.-H. Lee et al., 2010).

Taken together, our pharmacological data and RNAi results reveal differential effects of *ubh-4* and *usp-14* on the number of autophagosomes and autolysosomes, and a partial tissue variation, thus highlighting a complexity in the dynamics of autophagy in *C. elegans* (Fig 6).

## Discussion

In this study, we show that genetic downregulation or pharmacological impairment of the proteasome-associated DUBs Uchl5/UBH-4 and Usp14 reduces autophagy in human HeLa cells, and elicits differential effects on the pool size of APs and ALs in various tissues in *C. elegans.* Our study reveals that downregulation of *Uchl5* by siRNA causes both increased GFP/RFP ratio and accumulation of LC3-II and p62 in GFP-LC3-RFP-LC3ΔG HeLa cells, suggesting a reduction in autophagy (Fig 1). Similarly, siRNA knockdown of *Usp14* increases the GFP/RFP ratio and amount of LC3-II and p62, reflecting perturbed autophagy (Fig S3). It is worth noticing that HeLa cells have been reported to contain high levels of LC3-II (Tanida et al., 2005), and that although LC3-II is the commonly used marker for autophagosomes, interpreting the outcome of LC3-II levels on the autophagy process is not straight forward (Klionsky et al., 2021). For example, increased LC3-II levels could be due faster conversion from LC3-I or block in autophagosome-lysosome fusion step (Klionsky et al., 2021). As we detected accumulation of LC3-II together with p62, our results show slower degradation of autophagosomal substrates indicative of reduced autophagy. In agreement with our study, Kim et al. (E. Kim et al., 2018) reported reduced autophagic flux, i.e., increased levels of both LC3-II and p62 in HEK293 cells and MEF cells upon *Usp14* downregulation. In contrast, downregulation of *Usp14* has been shown to enhance autophagy in human H4 neuroglioma cells, as increased LC3-II levels concomitant with reduced p62 levels were detected (D. Xu et al., 2016).

As a complementary approach to the genetic downregulation of *Uchl5* and *Usp14*, we investigated the effect of the commonly used DUB inhibitors b-AP15, a dual inhibitor of Uchl5 and Usp14, and the Usp14-specific inhibitor IU-1 on autophagy (B.-H. Lee et al., 2010) (D’Arcy et al., 2011). D’Arcy and co-workers have shown that b-AP15 functions by blocking the deubiquitinating activity of Uchl5 and Usp14 in the 19S regulatory particle of the proteasome, thereby causing accumulation of proteasomal substrates *in vitro* and in human cell lines (D’Arcy et al., 2011). IU1 treatment on the other hand has been reported to decrease the ubiquitin chain trimming capacity of Usp14 leading to increased degradation of proteasomal substrates (B.-H. Lee et al., 2010). Currently, there is no commercially available inhibitor specifically targeting Uchl5, and therefore, we used comparison studies between b-AP15 and IU1 treatments to analyze the role of Uchl5 on autophagy. Our data show that both b-AP15 treatment and IU1 treatment result in reduced autophagy, as increased GFP/RFP ratio as well as accumulation of LC3-II and p62 were observed in both cases (Fig 2 and S3). Thus, Uchl5 and Usp14 appear to have a similar modulatory effect on autophagy in GFP-LC3-RFP-LC3ΔG HeLa cells. It is unlikely that the impact of Usp14 on autophagy would override that of Uchl5, as our RNAi results show that both Usp14 and Uchl5 knockdown reduces autophagy. Supportive results on LC3-II accumulation upon b-AP15 treatment have previously been shown in triple negative breast cancer (TNBC) cell lines (Vogel et al., 2015). It has also been reported that the DUB inhibitor NiPT, which also blocks both Uchl5 and Usp14, induces autophagy in A549 and NCI-H1299 lung cancer cell lines (J. Chen et al., 2020). In agreement with a previous study by Kim et al. (E. Kim et al., 2018), we demonstrate that IU1 treatment impairs autophagy, as detected by increased GFP/RFP ratio as well as accumulation of LC3-II and p62. A more complex view on the effect of IU1 treatment on autophagy is described by Xu and co-workers (L. Xu et al., 2020) by showing that lower concentration increases the levels of both LC3-II and p62, whereas higher concentration increases LC3-II but decreases p62 levels in HeLa cells. In our case, we detected increased levels of both LC3-II and p62 upon treatment with a similar high concentration of IU1, which could be due to different treatment duration and/or the transgenic GFP-LC3-RFP-LC3ΔG HeLa cell line. A study by Srinivasan and co-workers reveals that IU1 treatment differentially influences autophagy flux i.e., the levels of LC3-II upon blockage of fusion of autophagsomes with lysosomes, in ML1 and primary thyroid cells, but without a concurrent effect on p62 levels (Srinivasan et al., 2023).

Our results extend further support for a functional interplay between UPS and ALP, where impairment or activation of proteasome has previously been reported to cause induction or reduction of autophagy, respectively. For instance, pharmacological inhibition of the proteasome by lactacystin enhances autophagy in SH-SY5Y neuroblastoma cells and in a UPS-compromised mice model (Shen et al., 2013). Similarly, treatment with the MG-132 proteasome inhibitor activates autophagy in rat alveolar macrophage cells (Fan et al., 2016) and in HEK293 cells (Li et al., 2019). Pharmacologically or genetically induced impairment of the proteasome enhances autophagy in the cardiomyocytes of mice (Q. Zheng et al., 2011) (Pan et al., 2020), and RNAi targeting of different proteasome subunits results in both enhanced basal autophagy as well as starvation-induced autophagy in drosophila larvae (Lőw et al., 2013). Conversely, stimulation of proteasomal substrate degradation through downregulation of Usp14 results in reduced autophagic flux in HEK293 and MEF cells (E. Kim et al., 2018). Further, increased proteasome levels and activity correlate with reduced autophagy in the retina of rhodopsin P23H mutant mice treated with the phosphodiesterase-4 inhibitor rolipram (Qiu et al., 2019). In this study, we show that modulation of the proteasome via genetic downregulation or pharmacological inhibition of the proteasome-associated DUBs Uchl5 or Usp14 causes reduced autophagy in human GFP-LC3-RFP-LC3ΔG HeLa cells (Figs 1, 2 and S3).

Our results also reveal how modulating UPS via the proteasome-associated DUBs Uchl5/UBH-4 and USP-14 affect autophagy at the tissue level in *C. elegans.* Autophagy plays a key role in various developmental and physiological processes in *C. elegans* including embryogenesis, development, dauer formation, longevity, and stress responses (Meléndez et al., 2003) (Hansen et al., 2008) (Zhao et al., 2009) (Tian et al., 2010) (Alberti et al., 2010) (Wu et al., 2015) (Palmisano & Meléndez, 2019) (Y. Chen et al., 2021). In adult animals, autophagy has been previously investigated in intestinal cells, hypodermal seam cells, neurons, muscle cells, and the pharynx (Chapin et al., 2015) (Zhang et al., 2015) (Chang et al., 2017) (H. Zheng et al., 2020).

We show that downregulation of *ubh-4* or *usp-14* by RNAi has different effects on the pool size of AP and AL in intestinal cells, hypodermal-seam cells, and pharynx (Fig 3, 4 and 6). The *usp-14* RNAi appears to block autophagosome formation both in the intestinal and hypodermal seam cells without affecting the rate of autophagosome-lysosomal fusion and *ubh-4* knockdown has a similar effect in intestinal cells. However, in hypodermal seam cells *ubh-4* downregulation leads to increased number of autolysosome without a change in autophagosome number, suggesting that *ubh-4* downregulation slowers the rate of autophagy flux in this tissue. In agreement with our tissue differential effect of DUBs on autophagy, several studies have reported that autophagy varies in different tissues in *C. elegans.* According to Chapin et al. there are cell type-specific differences in the overall rate of autophagic flux both in basal and stress-induced autophagy in *C. elegans* (Chapin et al., 2015), and tissue variations in the age-dependent decrease in autophagy have been reported (Chang et al., 2017). Zheng et al have revealed that the autophagy genes are involved in a tissue- and stage-specific manner during the development of *C. elegans* (H. Zheng et al., 2020). Additionally, dietary restriction has been shown to result in reduction of the number of autophagosomes in intestinal cells (Gelino et al., 2016).

Compared to the RNAi-induced autophagy response in the intestine and hypodermal seam cells, the same animals did not display any phenotype in the pharynx after the three days of RNAi treatment, not even when targeting *lgg-1* or *rab-7* (Fig S5A and S5B). However, continuous exposure of the animals to *ubh-4* or *usp-14* RNAi treatments for two generations resulted in a decreased number of autophagosomes and autolysosomes (Fig 4B), suggesting a reduced autophagic flux. Previous studies have also reported that pharynx is resistant to first generation RNAi, but sensitive in animals upon continuous RNAi feeding for two generations (Kumsta & Hansen, 2012) (Shiu & Hunter, 2017). Downregulation by RNAi has its limitations, but in comparison to a potential compensatory redundancy and a chronic effect in knockout animals, RNAi and/or pharmacological inhibition studies offer a way to mimic a stress or disease condition.

In agreement with our RNAi studies targeting *ubh-4* and *usp-14,* pharmacological inhibition of these DUBs induced variation in the pool size of autophagosomes and autolysosomes in different tissues in *C. elegans* (Fig 5). For validation study, we tested these inhibitors on a previously established polyubiquitin reporter strain and observed that b-AP15 treatment leads to increased fluorescent indicating enhanced accumulation of polyubiquitinated proteins (Fig S6A and S6B), whereas animals treated with IU1 caused decreased amount of polyubiquitinated proteins in the intestine. These results are in agreement with the human cell line data, where accumulation of proteasomal substrates upon b-AP15 treatment and increased degradation of proteasomal substrates upon IU1 treatment have been shown (B.-H. Lee et al., 2010) (D’Arcy et al., 2011). Our b-AP15 results on polyubiquitin accumulation differ from our previously published result on the effect of *ubh-4* RNAi which showed that downregulation of *ubh-4* results in decreased accumulation of the polyubiquitin reporter, and increased proteasome activity *in vivo* and *in vitro* (Matilainen et al., 2013). The difference between the b-AP15 and *ubh-4* RNAi effect is likely due to the dual inhibitory effect of b-AP15 on UBH-4 and USP-14.

Monitoring autophagy in an adult multicellular organism is intricate due to the diverse and morphologically distinct cell types with spatial and temporal differences in responses to physiological or pathophysiological stress conditions. Altogether, our data reveal that modulation of UPS via pharmacological or genetic impairment of the proteasome-associated DUBs UBH-4 and Usp14 differentially affects autophagy in a tissue variable manner. However, future studies are required to reveal the molecular mechanism underlying the tissue-specific differences in autophagy in response to Uchl5/UBH-4 and Usp14. Our results thus expand the existing knowledge on the wide variety in the kinetics and regulation of autophagy in different cell types (Klionsky et al., 2021) and highlight the dynamic complexity of autophagy. This tissue/cell type specific information is valuable for future development of therapeutic options targeting autophagy or autophagy-associated diseases.

## Material and Methods

### Mammalian cell cultures

HeLa cells expressing GFP-LC3-RFP-LC3ΔG were cultured in high glucose DMEM. The media was supplemented with 10% fetal bovine serum (FBS), L-glutamine, penicillin and streptomycin and maintained in a 5% CO_2_ incubator. For siRNA experiments, FlexiTube GeneSolution for *uchl5 and usp14* (QIAGEN) and AllStars Negative Control siRNA (QIAGEN) were used with HiPerFect transfection Reagent (QIAGEN). Cells treated with siRNA were incubated for 48hr prior to live imaging or sample collection for quantitative analysis. For pharmacological treatment, we used 1 uM of b-AP15 (Sigma-Aldrich, Cat# 662140) or 100 uM IU-1 (Sigma-Aldrich, Cat# 662210). Cells were treated with b-AP15 for 18h and IU-1 for 6 h. DMSO (Fisher Scientific, Cat# 15498089) is used as control.

### Microscopy, equipment, and image analysis

GFP-LC3-RFP-LC3ΔG expressing HeLa cells transfected with *uchl5* or *usp14* siRNA or treated with inhibitors were grown in Thermos Scientific^TM^ Nunc^TM^ Lab-Tek^TM^ II chambered coverglass plates (Fisher scientific, cat# 16260671) and imaged after 48 hr. Live cells were imaged with a Zeiss LSM880 confocal microscope (Motorized Zeiss Axio Observer.Z1 inverted microscope), at 63x 1.4 NA plan-Apochromat objective and at 37°C and 5% CO_2_. Confocal images were converted to tiff-format using Zen 2 lite (blue). Images were quantified from the original version without any modification, using Fiji ImageJ software. The number of puncta was counted using plugins, FeatureJ, and FeatureJ Laplacian commands from Fiji software. A threshold was selected so that most of the dots were selected for all images, and then the analyze particle command gave the number of puncta present on that image. For quantification of the total fluorescence intensity of images, we used Fiji ImageJ software. The background was subtracted using the corresponding command in Fiji software. Threshold was selected for the brightest images and the same threshold was applied to all images from the same experiment. The average of mean intensity was analyzed.

All the images were processed with Fiji ImageJ software. The brightness of the images was increased in the same way to all corresponding images from the same experiment, to be able to make the fluorescent signal clearly visible.

### Western blotting

For Western blotting, cell lysates were collected after 48hr of siRNA treatment or 18hr of b-AP15 or 6hr of IU-1 treatment. The cell lysates were lysed by vigorous vortexing. Samples were run on SDS-PAGE gel and immunoblotted onto a nitrocellulose membrane using Trans-Blot Turbo transfer system (Bio-Rad). Anti-LC3B antibody (anti-rabbit, Sigma-Aldrich, Cat# L7543, 1:5000 dilution), anti-P62 antibody (anti-rabbit, Sigma, Cat# P0067, 1:10,000 dilution), anti-UCHL5 (anti-mouse, Santa Cruz, cat# sc271002, 1:500 dilution), anti-USP14 (anti-mouse, Sigma, Cat#SAB1406778, 1:2500 dilution) and anti-HSC70 antibody (anti-mouse, Santa Cruz, Cat# sc7298, 1:5000 dilution) were used for immunoblotting. For the anti-LC3B antibody and the anti-P62 antibofy, the secondary antibody used was anti-rabbit IgG-HRP conjugate (LA_W4011,1:10,000 dilution), and for anti-UCHL5, anti-USP14 and HSC70 anti-mouse IgM-HRP conjugate (Calbiochem, Cat# 401225,1:10,000 dilution). Image Studio software (Licor) was used for imaging and quantifying the signals.

### Quantitative real-time PCR

Cell lysates were collected after 48hr of siRNA treatment and stored at -80°C. Total RNA was extracted from the freezed samples using NucleoSpin RNA kit (Macherey-Nagel) and concentration of extracted RNA was measured with Nanodrop spectrophotometer at 260 nm. RT-PCR was performed using Maxima First Standard cDNA Synthesis Kit for RT-qPCR (Fermentas). The quantitative real-time PCR was done using Maxima SYBR Green/ROX qPCR Master Mix (2X) (Fermentas) and LightCycler 480 (Roche) quantitative PCR machine. The data from qPCR were normalized to the geometric mean of mRNA concentration of two reference genes (*gapdh* and *cyclophilin*).

### *C. elegans* and growth conditions

*C. elegans* strains were cultured and maintained as described previously (Brenner, 1974) under standard conditions at 20°C, on nematode growth medium (NGM) plates seeded with OP50. N2 (Bristol) and MAH215 were obtained from the Caenorhabditis Genetics Center (CGC).

### *C. elegans* RNA interference (RNAi)

RNAi was performed using the feeding method as described earlier (Timmons et al., 2001). The HT115 bacterial strain carrying the empty *pL4440* expression vector was used as a control. RNAi clones used in this study were from J. Ahringer library. The double stranded RNA expression was induced by adding 0.4 mM of isopropyl-β-D-thiogalactopyranoside (IPTG) (I6758, Sigma) and its concentration was further increased to 0.8 mM prior to seeding the plates. Unless otherwise indicated, age-synchronized animals were placed on control as well as RNAi seeded plates targeting *ubh-4* and *usp-14* as L1 larvae (day 1). For imaging experiments, *ubh-4* and *usp-14* RNAi treated animals were imaged at first day of adulthood (day 4).

### Inhibitor treatment

Chemicals were added to the nematode growth medium (NGM) prior pouring to the plates. We used DMSO (Fisher Scientific, Cat# 15498089) as a control, 10 uM of b-AP15 (Sigma-Aldrich, Cat# 662140) or 100 uM IU-1 (Sigma-Aldrich, Cat# 662210). NGM plates with chemicals were seeded with OP50 Escherichia coli bacteria. Once the plates were dried, age synchronized animals were placed on chemical seeded plates on the same day as L1 larvae (Day 1). Chemical treated animals were imaged at first day of adulthood (day 4).

### Microscopy of *C. elegans* and quantitative image analysis

Age synchronized animals were imaged at the first day of adulthood (Day 4). Animals were mounted on 3% agarose pad on glass slides and immobilized using 1 mM levamisole in M9 buffer (22 mM KH_2_PO_4_, 41 mM Na_2_HPO_4_, 8,5 mM NaCl and 19 mM NH_4_Cl). Autophagy dual marker reporter strains were imaged using LSM780 confocal microscope (Motorized Zeiss Axio Observer.Z1 inverted microscope), z-stack images were acquired at 0.8 µm slice intervals at 40x 1.3 NA plan Neofluor objective. Z-stack images were converted to maximum intensity projection format using ZEN 2.1 (black) and converted to tiff-format using Zen 2 lite (blue). The number of puncta was calculated manually.

All the images were processed with Fiji ImageJ software. The brightness of the images was increased in the same way to all corresponding images from the same experiment, to be able to make the fluorescent signal clearly visible.

### Quantitative real-time PCR

Age synchronized RNAi treated animals were collected in M9 at first day of adulthood (day 4) and stored at -80°C. Total RNA was extracted from the freezed samples using NucleoSpin RNA kit (Macherey-Nagel) and concentration of extracted RNA was measured with Nanodrop spectrophotometer at 260 nm. RT-PCR was performed using Maxima First Standard cDNA Synthesis Kit for RT-qPCR (Fermentas). The quantitative real-time PCR was done using Maxima SYBR Green/ROX qPCR Master Mix (2X) (Fermentas) and LightCycler 480 (Roche) quantitative PCR machine. The data from qPCR were normalized to the geometric mean of mRNA concentration of three reference genes (*act-1, cdc-42* and *pmp-3)* (Vandesompele et al., 2002).

### Statistical analysis

Statistical significance was determined using the student’s t-test (two-tailed).

## Acknowledgments

We thank the Biomedicum Imaging Unit (BIU), Faculty of Medicine, University of Helsinki, for their help with microscopy and image analysis, Holmberg lab member Elisa Mikkonen for her comments on the figures. Some strains were provided by the CGC, which is funded by NIH Office of Research Infrastructure Programs (P40 OD010440).

## Competing interests

The authors declare no conflict of interest.

## Funding

This study was supported by grants to CIH from the Academy of Finland (297776), Sigrid Jusélius Foundation and the Medicinska Understödsföreningen Liv och Hälsa r.f. S.J. was supported by the Doctoral Programme in Biomedicine, University of Helsinki and grants from the Magnus Ehrnrooth Foundation and the Finnish Cultural Foundation.

## Data availability

The data and tools that support the findings of this study are available from the corresponding author upon request.

## Author contribution

Conceptualization: S.J., C.I.H.; Methodology: S.J., C.I.H.; Software: S.J.; Validation: S.J., J.P., C.I.H.; Formal analysis: S.J.; Investigation: S.J., J.P.; Resources: C.I.H.; Data curation: S.J., C.I.H.; Writing - original draft: S.J.; Writing - review & editing: S.J., J.P., C.I.H.; Visualization: S.J., C.I.H.; Supervision: C.I.H.; Project administration: C.I.H.; Funding acquisition: S.J., C.I.H.

---

**Figure S1.**
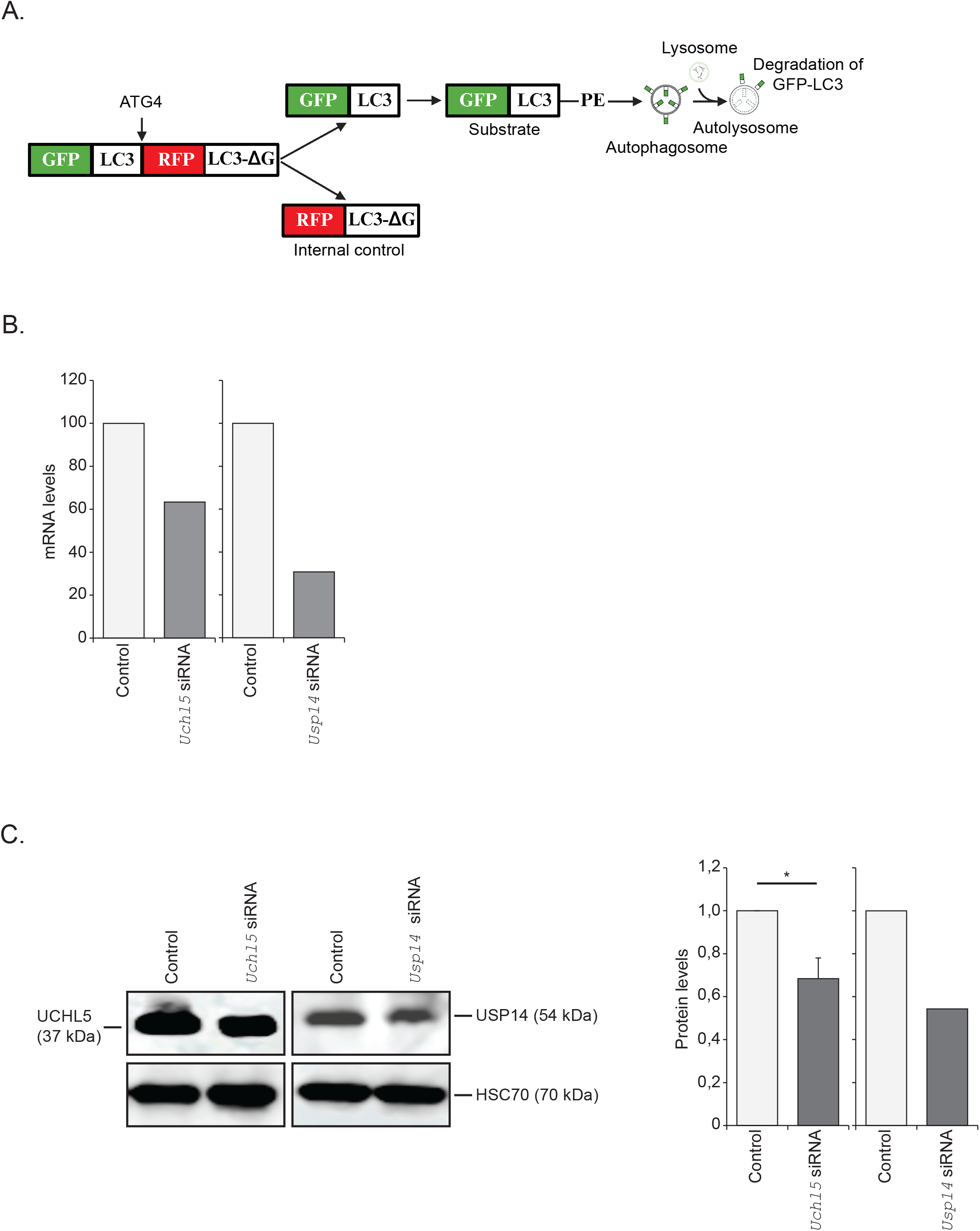
Downregulation of *Uchl*5 and *Usp14* upon treatment with siRNA. **A.** Schematic representation of the GFP-LC3-RFP-LC3ΔG fluorescent probe (Kaizuka et al., 2016). **B.** GFP-LC3-RFP-LC3ΔG HeLa cells treated with control, *Uchl5* or *USsp14* siRNA and collected 48h post-transfection. Expression of *Uchl5* or *Usp14* mRNA was measured with qPCR. Graph shows the percentage change in the mRNA levels compared to control (set as 100%). **C.** Whole cell extracts (48h post-transfection) were analyzed by SDS-PAGE and immunoblotted against Uchl5, Usp14 and HSC70. The graphs (on right panel) show average fold change in levels of Uchl5 and Usp14 normalized against HSC70. Results are the mean of quantifications from three (Uchl5) or one (Usp14) independent experiment. Error bars, SEM, *p<0,05 compared to the control (set as 1).

**Figure S2.**
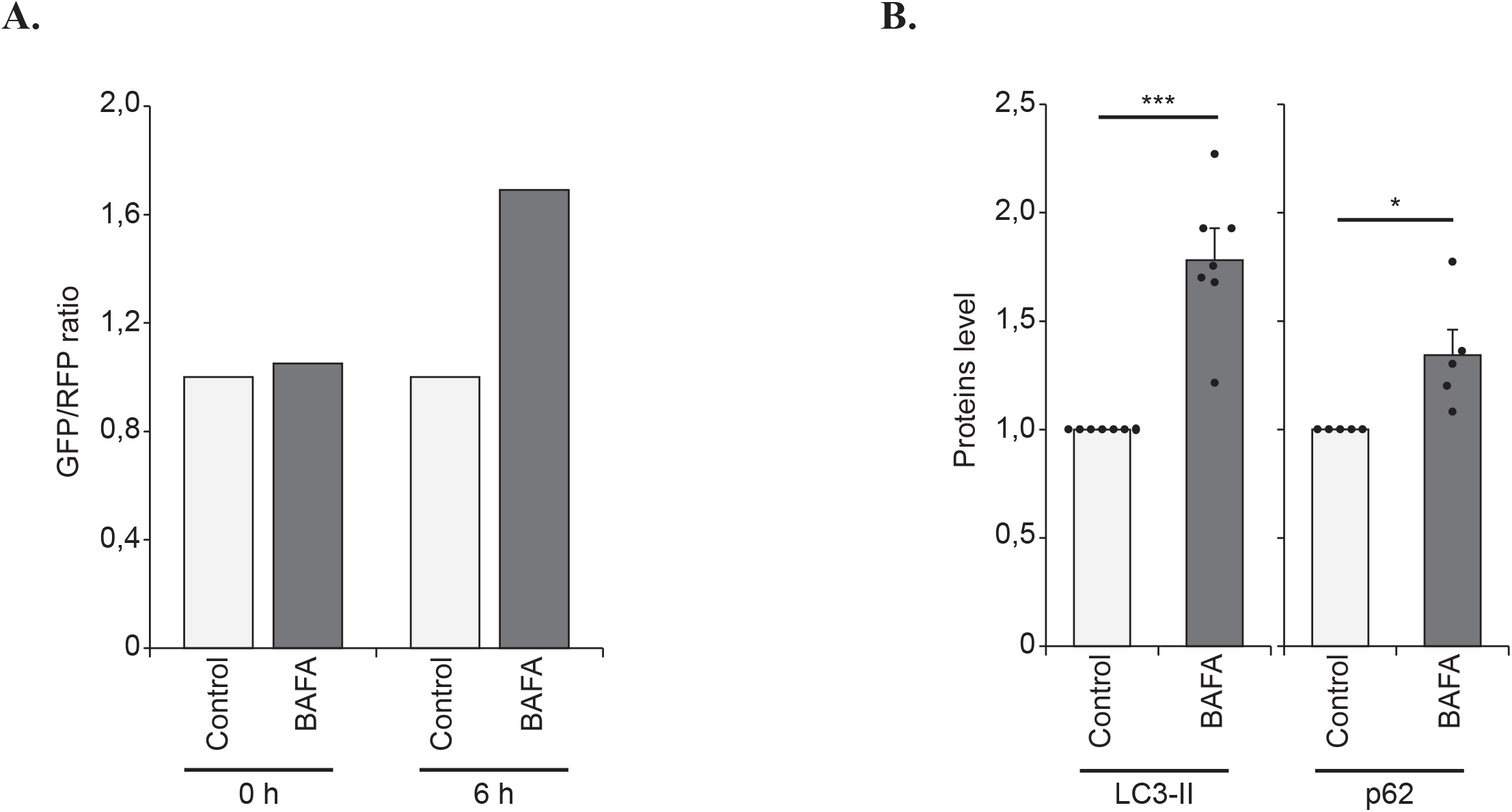
Validation experiments using BAFA. **A.** GFP-LC3-RFP-LC3ΔG HeLa cells treated with control (DMSO) or BAFA (100 nM) for 6h. The graph shows the quantification of the relative fold change in the ratio of GFP/RFP per image (Control set at 1) (Five images were analyzed). **B.** GFP-LC3-RFP-LC3ΔG HeLa cells treated with control (DMSO) or BAFA (100 nM) for 6h. Whole cell lysates were analyzed by SDS-PAGE and immunoblotted against LC3, p62 and HSC70. The graphs show average fold change in levels of LC3 and p62 normalized against HSC70. Results are the mean of quantifications from three independent experiments. Error bars, SEM, *p<0,05, ***p<0,001 compared to the control (set as 1).

**Figure S3.**
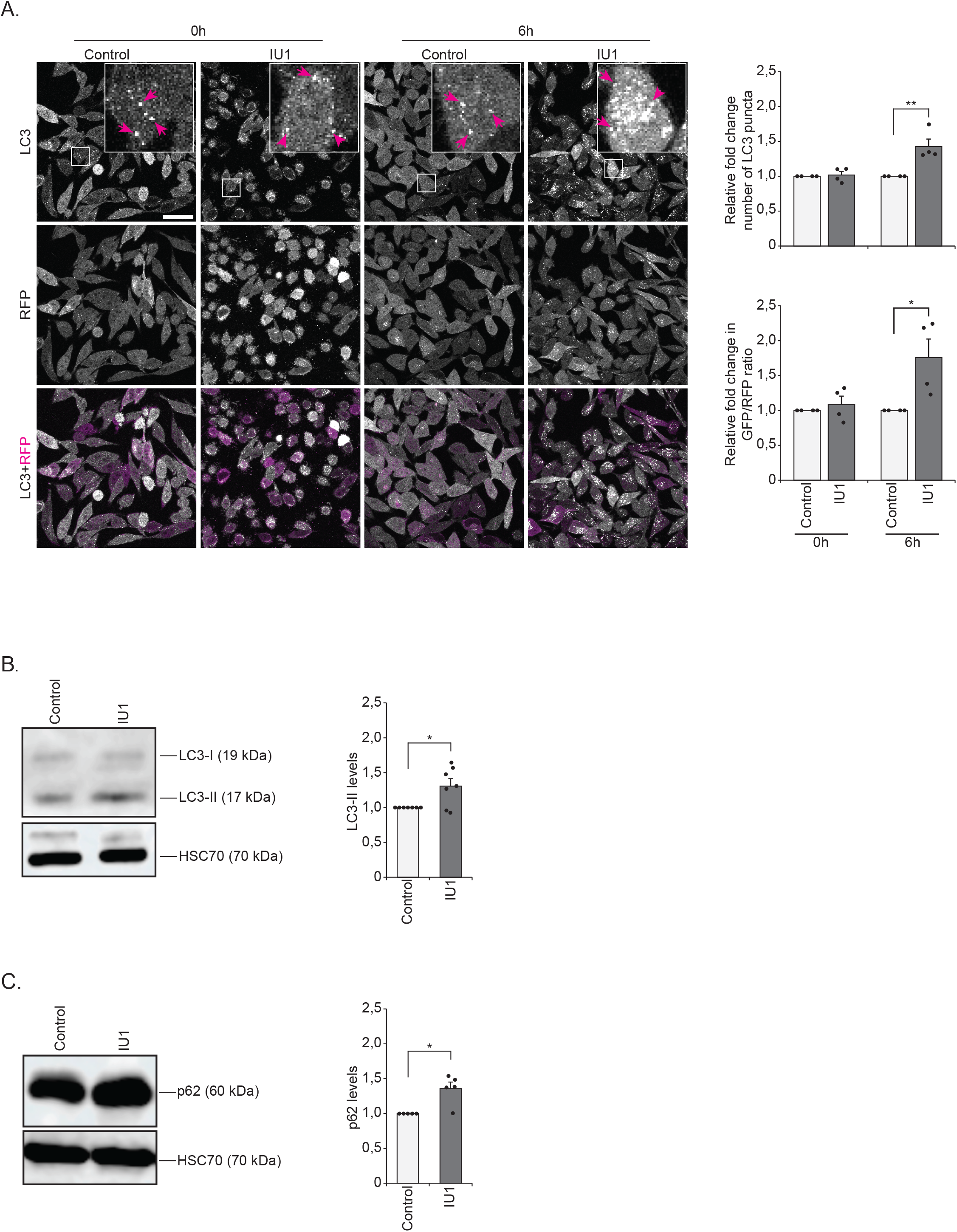
Pharmacological inhibition of Usp14 using IU1 inhibitor reduces autophagy. **A.** Fluorescence confocal images of control (DMSO) or IU1 (100µM) treated GFP-LC3-RFP-LC3ΔG HeLa cells after 6h post-treatment. Insets show enlarged view of the indicated areas. Magenta arrows point to some of the puncta. Scale bar, 20 µm. The right upper graph shows the quantification of relative fold change in the number of GFP-LC3 puncta per image (Control set at 1). The right lower graph shows the quantification of the relative fold change in the ratio of GFP to RFP per image (Control set at 1). Results are from four independent experiments (total 15-20 images were analyzed). Error bars, SEM, *p<0,05, **p<0,01 compared to control. **B and C**. GFP-LC3-RFP-LC3ΔG HeLa cells treated with control or IU1 (100µM) for 6h. Whole cell lysates were analyzed by SDS-PAGE and immunoblotted against LC3, p62 and HSC70. The graphs (on right panel) show average fold change in levels of LC3 (B) and p62 (C) normalized against HSC70. Results are the mean of quantifications from six independent experiments. Error bars, SEM, *p<0,05 compared to the control (set as 1).

**Figure S4.**
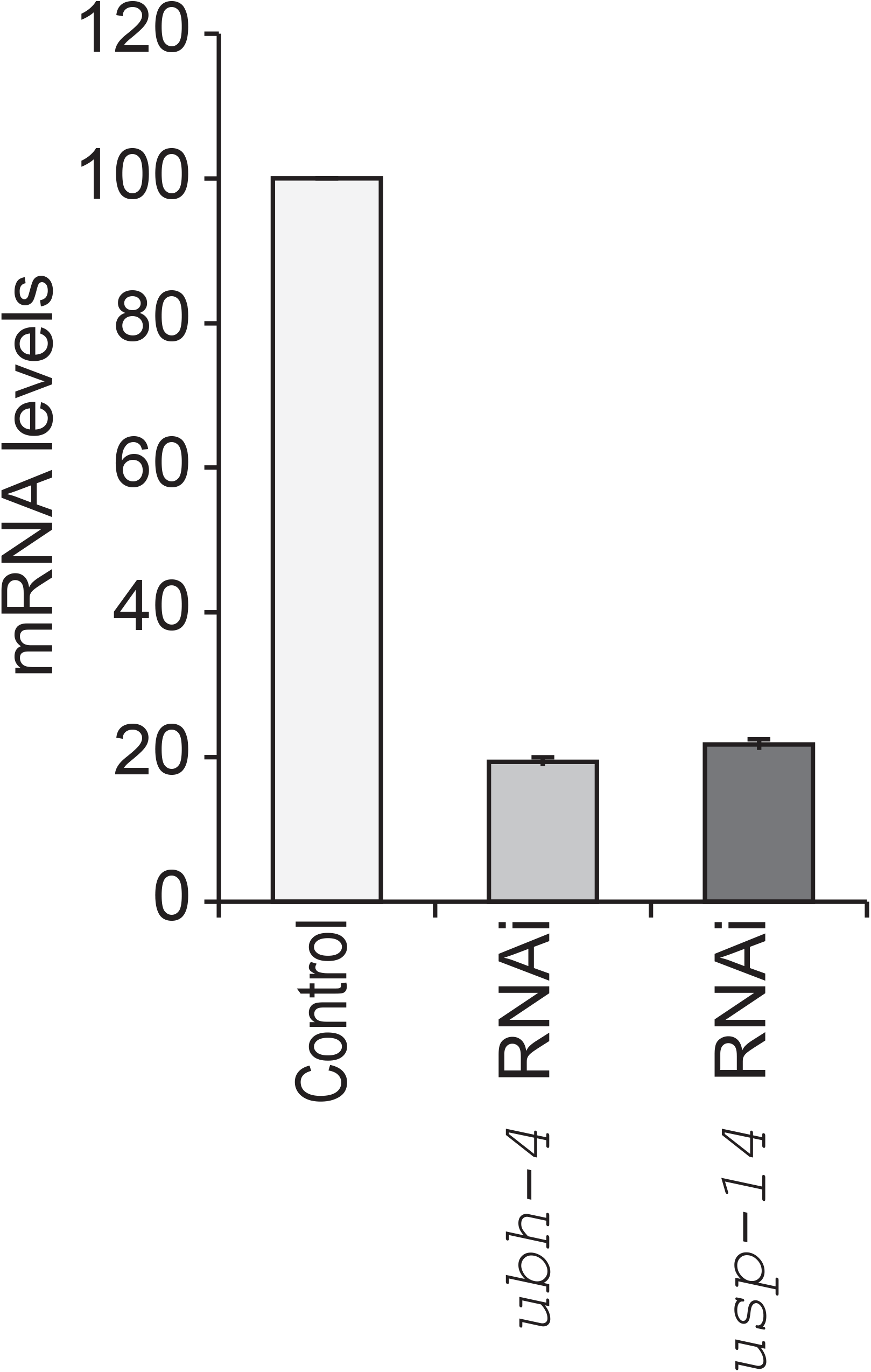
Efficient downregulation of *ubh-4* and *usp-14* upon RNAi. Wild-type animals were exposed to control, *ubh-4* or *usp-14* RNAi treatment starting at the L1 larval stage and collected at Day 1 of adulthood. Expression of *ubh-4* or *usp-14* mRNA in the RNAi-treated animals was checked with qPCR. Graph shows percentage change in mRNA levels compared to control (set as 100%). The results are the mean of two independent experiments in triplicate. Error bar, STD.

**Figure S5.**
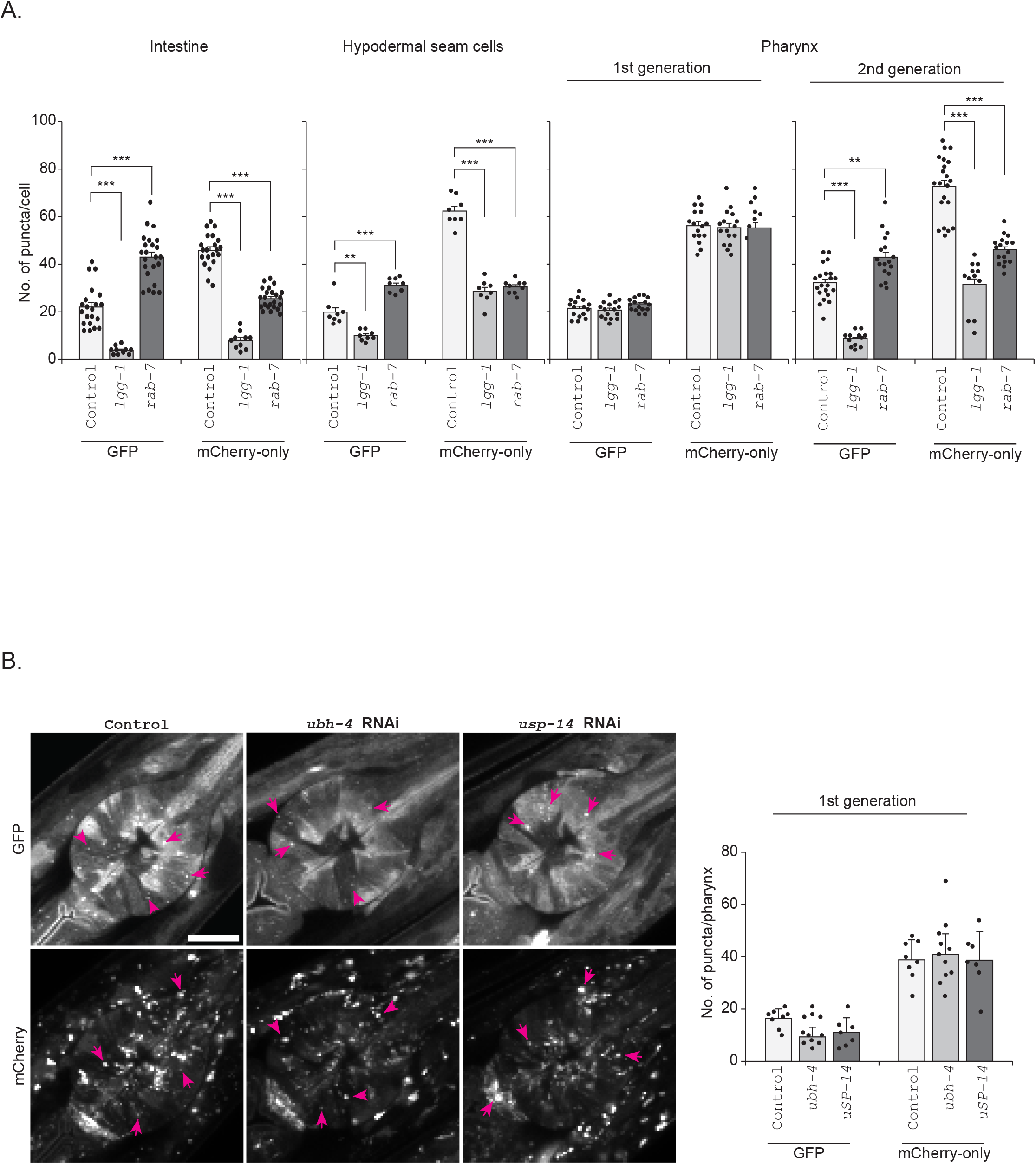
Effect of downregulation of autophagy genes or proteasome-associated DUBs on autophagy in different tissues. **A** Graphs show the quantification of the number of puncta positive for GFP and mCherry-only in the intestinal cells, hypodermal seam cells and the pharynx. All animals, except the ones labelled second generation, were exposed to RNAi from L1 larval stage to day 1 of adulthood (First generation). The animals and their progeny were further continuously exposed to RNAi, and the day 1 of adulthood of the progeny is here labelled as second generation. Results are from three independent experiments. Puncta were counted from a total of 20-30 individual corresponding cells. Error bars, SEM, **p<0,01, ***p<0,001 compared to control. **B.** Fluorescence confocal micrographs of control, *ubh-4* or *usp14* RNAi-treated mCherry::GFP::LGG-1 animals showing the pharynx at day 1 of adulthood (First generation). Graphs show the quantification of the number of puncta positive for GFP and mCherry-only. Results are from three independent experiments. Puncta were counted from a total of 10-12 pharynges from 10-12 animals. Error bars, SEM.

**Figure S6.**
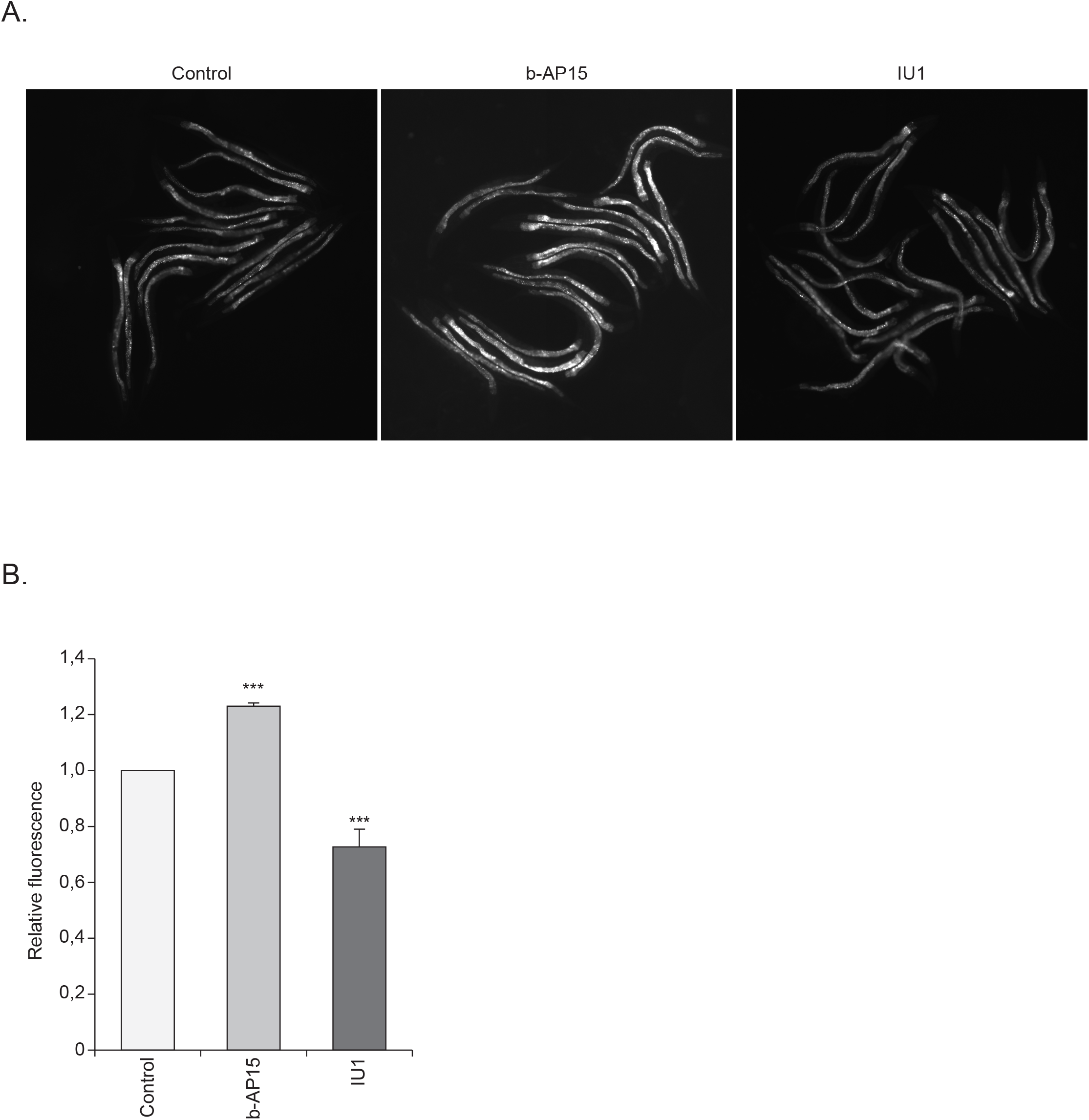
The b-AP15 and IU1, inhibitors of the proteasome-associated DUBs UBH-4 and USP-14, affect accumulation of polyubiquitinated proteins in intestinal cells. **A.** Representative fluorescence micrographs of control (DMSO), b-AP15 (10 µM) or IU1 (100 µM) treated animals expressing the polyubiquitin reporter in the intestinal cells. Animals were treated with the inhibitors from L1 larval stage till day 1 of adulthood. **B.** Graph shows quantification of fluorescence, which reflects accumulation of polyubiquitinated proteins in the intestinal cells. Results are from three independent experiments (number of animals 90-100). Error bars, SEM, ***p<0,001 compared to control.

## Notes

### Competing Interest Statement

The authors have declared no competing interest.

